# Chromosome-scale nuclear and mitochondrial genome assemblies reveal extreme genome compartmentalization in the hazelnut pathogen *Anisogramma anomala*

**DOI:** 10.64898/2026.10.20.760510

**Authors:** Alex Z. Zaccaron, Kelsey L. Søndreli, Jared M. LeBoldus, Alexandra J. Weisberg

## Abstract

*Anisogramma anomala* is an obligate biotrophic ascomycete and the causal agent of Eastern filbert blight, the most economically important disease of hazelnut in the United States. Here, we generated a chromosome-scale reference genome for *A. anomala* isolate OR-Ennis-d and characterized its nuclear and mitochondrial genome architecture. The 344-Mb nuclear genome was resolved into 9 predicted chromosomes and is repeat-rich, with transposable elements comprising 88.8% of the assembly. Although transposable elements are abundant, their divergence landscape and low retention of conserved protein domains support a history of past extensive transposon proliferation followed by widespread degeneration. Repeat accumulation was highly uneven and generated pronounced genome compartmentalization. Approximately half of predicted genes occur in compact, TE-poor regions with short flanking intergenic regions that are enriched for conserved BUSCO orthologs. In contrast, genes in TE-rich, gene-sparse regions with long flanking intergenic regions are enriched for carbohydrate-active enzymes, secreted proteins, and candidate secreted effector proteins, consistent with a two-speed-like genome architecture. The 462-kb mitochondrial genome is among the largest reported for ascomycetes and contains 117 introns, including an intron adjacent to *cob* codon 143 that may prevent acquisition of the p.G143A substitution associated with resistance to quinone outside inhibitor fungicides. Finally, we identified mitochondrial DNA insertions at 20 nuclear loci totaling at least 616.9 kb, an unusually extensive amount for a fungal genome. This chromosome-scale genome resource reveals the extreme nuclear and mitochondrial genome architecture in *A. anomala* and provides a foundation for investigating genome evolution and adaptation to hazelnut resistance.

## Introduction

*Anisogramma anomala* (Peck) E. Müller (Ascomycota, Gnomoniaceae) is an obligate biotrophic pathogen that causes Eastern filbert blight (EFB) of hazelnut (Corylus spp.) (Gottwald and Cameron 1979; EFSA Panel on Plant Health (PLH) et al. 2018). In the USA, hazelnuts represent a major specialty crop in the Pacific Northwest (PNW), particularly in Oregon’s Willamette Valley, where approximately 98% of the domestic hazelnut supply is produced (Julian et al. 2009). EFB is considered the main limiting factor for hazelnut production in the USA, and effective management of this disease is tightly linked with sustainability and profitability of the hazelnut industry (Thompson et al. 1996).

Native to eastern North America, *A. anomala* co-evolved with the American hazelnut (*Corylus americana*), a host that exhibits substantial tolerance to EFB (Gottwald and Cameron 1979; Pinkerton et al. 1995; Revord et al. 2020). In contrast, the commercially important European hazelnut (*C. avellana*) is highly susceptible to EFB. Historically, commercial production of European hazelnut in the PNW occurred outside the native range of *A. anomala*. However, the pathogen was accidentally introduced into southwestern Washington during the late 1950s, likely through infected plant material, and subsequently spread throughout the PNW (Davison and Davidson 1973; Cameron 1976; Pscheidt et al. 2022; Tobia et al. 2024). In response, the breeding programs at Oregon State University played a central role in mitigating EFB through the development of resistant cultivars (Mehlenbacher et al. 2007; Mehlenbacher et al. 2009; Mehlenbacher et al. 2011; Mehlenbacher et al. 2013; Mehlenbacher et al. 2014). A major development came with the deployment of the ‘Gasaway’ resistance locus, which confers resistance to EFB and has been widely incorporated into commercial hazelnut varieties (Mehlenbacher et al. 1991). For decades, this resistance was effective and became a milestone of EFB management in the PNW. However, the extensive reliance on a single resistance source has likely imposed strong selection pressure on the *A. anomala* population. Since 2022, isolates capable of overcoming ‘Gasaway’-mediated resistance have emerged, raising concerns about the durability of resistance and the long-term stability of hazelnut production systems (Wiman 2023). In many fungal pathogens, mutations in effectors or other pathogenicity-related genes can enable pathogens to evade recognition by host resistance mechanisms. Single-nucleotide substitutions, gene deletions, or altered expression of effector genes can restore virulence in previously avirulent strains (Fudal et al. 2007; Mesarich et al. 2023). Larger-scale genomic structural variations, such as large deletions, translocations, and duplications can also alter pathogenicity or virulence, and are often facilitated by the presence of transposable elements (TEs) (Chuma et al. 2011; Manning et al. 2013; Zaccaron and Stergiopoulos 2024).

Several fungal plant pathogens exhibit compartmentalized genomes in which gene-dense regions enriched in conserved genes coexist with repeat-rich, gene-sparse regions that may contain lineage-specific and host-interaction genes, including effectors (Dong et al. 2015; Frantzeskakis et al. 2019; Torres et al. 2020). These dynamic regions are frequently enriched in TEs, which can promote local genome instability through ectopic recombination, TE insertion, duplication, deletion, and chromosomal rearrangement (Seidl and Thomma 2017; Torres et al. 2020). In many ascomycete fungi, TE-rich regions are also subject to repeat-induced point mutation (RIP), a genome-defense process that introduces transition mutations into duplicated sequences and can extend into adjacent genomic regions (Galagan and Selker 2004; Rouxel et al. 2011). Consequently, pathogenicity-related genes located in repeat-rich compartments may experience elevated rates of sequence diversification, altered expression, copy-number variation, and presence–absence polymorphism, thereby facilitating adaptation to new hosts and the evasion of host resistance genes.

Mitochondrial genomes are an additional component of fungal genome architecture that can inform pathogen evolution (Black et al. 2021). Fungal mitogenomes vary widely in size and structure, largely because of variation in intron content, mobile intronic elements, intergenic regions, and gene order (Aguileta et al. 2014; Sandor et al. 2018). Mitochondrial introns can substantially expand fungal mitogenomes and may influence the evolution of mitochondrial genes. This is especially relevant for the mitochondrial cytochrome *b* gene (*cob*), which encodes the target of quinone-outside inhibitor (QoI) fungicides (Bartlett et al. 2002). Introns adjacent to *cob* codon 143 can prevent acquisition of the p.G143A substitution, a common fungicide-resistance mechanism in fungal pathogens (Fernández-Ortuño et al. 2008). Mitochondrial DNA can also be transferred to the nuclear genome as nuclear mitochondrial DNA segments (NUMTs), linking mitochondrial and nuclear genome evolution (Hazkani-Covo et al. 2010). However, the impact of NUMTs on fitness of fungal pathogen remains understudied.

Despite advances in understanding the genomic basis of virulence and host adaptation in other fungal plant-pathogen systems, these processes remain poorly understood in *A. anomala*. Functional characterization of candidate pathogenicity genes is constrained by the difficulties in culturing *A. anomala in vitro* and by its prolonged 12-16-month latent period in hazelnut, which complicates pathogenicity assays (Gottwald and Cameron 1979; Stone et al. 1994). Genomic investigation has been further challenged by the exceptionally large and repeat-rich *A. anomala* genome. An initial short-read assembly generated primarily to develop microsatellite markers spanned approximately 333 Mb but was highly fragmented (Cai et al. 2013). More recently, an assembly of isolate OR1 based on short-read and mate-pair sequencing generated a 342.5-Mb genome in 108 scaffolds (Cohen et al. 2024). Approximately 89% of the OR1 assembly is comprised of TEs and other repetitive sequences, predominantly Copia-like long terminal repeat retrotransposons, whereas many predicted genes occurred in GC-rich regions. Although these foundational resources established the high repeat content of *A. anomala*, limited chromosome-scale resolution constrained analyses of its long-range genome organization, repeat-rich regions, and the physical relationship between TEs and candidate host-interaction genes.

Addressing these limitations requires a highly contiguous long-read-based reference genome capable of resolving repetitive regions and their genomic context. Therefore, the objectives of this study were to: (1) generate a chromosome-scale reference genome for *A. anomala*; (2) characterize its nuclear and mitochondrial genome architecture; and (3) identify genomic features that may influence genome evolution and adaptation to hazelnut resistance.

## Materials and methods

### Fungal isolate and DNA extraction

Cankered branches of hazelnut (*Corylus avellana*) cv. ‘Ennis’ were collected from an orchard in Eugene, OR in 2024 and kept frozen in a -80°C freezer until DNA extraction. DNA was extracted from ascospores collected from stromata from a single branch canker, following a previously published protocol for ascospores collection (Cai et al. 2013; Cohen et al. 2024). Specifically, cankered branches were thawed at room temperature and surface sterilized by submerging them in sterile H_2_O solutions with 10% bleach (8.25% sodium hypochlorite) for two minutes with shaking, then in 70% ethanol for 30 seconds with shaking. Branches were then left to hydrate for 30 minutes in sterile H_2_O. A sterile scalpel was used to cut the neck of perithecia to expose a white mass containing the ascospores, which were then forced out by applying pressure under or adjacent to the stroma with a scalpel. The ejected white mass was then collected into a sterile mortar and pestle, ground with 10 mL of sterile H_2_O, and filtered through a 40 µm Cell Strainer (Corning, NY, USA, catalog no. 431750) into a 50 mL falcon tube.

Ascospores were pelleted by centrifugation at maximum speed for 5 min, the supernatant was discarded, and pellets were frozen at -80°C prior to DNA extraction. Genomic DNA was extracted using the QIAGEN Genomic-tip 20/G kit (Qiagen, Hilden, Germany; catalog no. 10223) with Genomic DNA Buffer Set (catalog no. 19060), following the manufacturer’s protocol with minor modifications. Briefly, spores were directly resuspended in 8 mL lysis buffer containing 20 mM EDTA, 10 mM Tris–Cl (pH 7.9), 0.5 mg mL⁻¹ lysing enzymes from Trichoderma, 1% Triton X-100, 500 mM guanidine-HCl, and 200 mM NaCl, and incubated at 37°C for 2 h with gentle agitation. RNase A was added to a final concentration of 20 µg mL⁻¹ and incubated for 30 min at 37°C, followed by Proteinase K treatment (0.8 mg mL⁻¹) for 2 h at 50°C with gentle agitation. Lysates were clarified by centrifugation at 12,000–15,000 × g for 20 min, and the supernatants were gently applied to QBT-equilibrated Genomic-tip columns to avoid DNA shearing. Columns were washed with Buffer QC and DNA was eluted with prewarmed (50°C) Buffer QF. DNA was precipitated with 0.7 volumes of room-temperature isopropanol, pelleted by centrifugation at 15,000 × g for 20 min, washed with ice-cold 70% ethanol, and air-dried. Pellets were resuspended in TE buffer and stored at -80°C until sequencing, minimizing pipetting to preserve high molecular weight DNA. Qubit 4 (Thermo Fisher Scientific, Singapore, catalog no. Q33226) and Nanodrop One (Thermo Fisher Scientific, Madison, WI, catalog no. ND-ONE-W) instruments were used to measure DNA quantity and quality. DNA fragment size was estimated by visualizing on a 0.75% agarose gel and with an Agilent TapeStation 4200 (Agilent Technologies, Santa Clara, CA).

### DNA sequencing

To prepare samples for Oxford Nanopore Technologies (ONT) sequencing, extracted DNA was first enriched for high molecular weight (HMW) fragments. To do so, a size selection step was performed using a 0.4× ratio of SPRI bead volume to DNA sample volume. Briefly, DNA elutions were brought up to 200 µl with ultrapure, molecular-grade water. After that, 80 µl of Ampure XP beads was added and gently resuspended into the diluted DNA, then incubated for 10 min at room temperature on a hula mixer. After incubation, a magnetic rack was used to pellet the beads, followed by three washes of 1 ml of 80% ethanol. The pellets were allowed to air-dry and were then resuspended in 30 µl of ultrapure water and incubated in a thermomixer shaker at 40°C and 300 rpm for 2 hours. After the beads were pelleted again, the DNA was eluted in nuclease-free water and quantified using a Qubit 4 DNA broad range assay kit. The Native Barcoding Kit 24 V14 (SQK-RBK114.24; ONT, Oxford, UK) was used to prepare long-read ONT libraries following the default protocol. The library was sequenced on a PromethION 2 Solo device (ONT, Oxford, UK) using an R10.4.1 flow cell and the ONT MinKnow software v23.07.5. Dorado v0.7.3 was used to perform basecalling using the dna_r10.4.1_e8.2_400bps_sup@v5.0.0 model (https://github.com/nanoporetech/dorado/). HMW DNA was also submitted to the University of Oregon Genomics and Cell Characterization Core Facility (Eugene, OR, USA) for Illumina sequencing. Libraries were prepared using the Watchmaker WGS Reduced Representation PCR-free workflow, following the manufacturer’s protocols. Sequencing was performed on an Illumina NovaSeq 6000 platform (Illumina, San Diego, CA, USA) using an S4 flow cell to generate 150 bp paired-end reads.

### Genome assembly

NanoPlot v1.43.0 and Filtlong v0.3.1 were used to perform quality control of nanopore reads (De Coster and Rademakers 2023; Wick 2025). Flye v2.9.4-b1799 with the parameter “--nano-hq” was used to generate a preliminary assembly from nanopore reads (Kolmogorov et al. 2019). The Foreign Contamination Screen (FCS) v0.5.4 tool with the fcs-gx database v2024-11-15 was used to identify assembled contigs representing bacterial contamination (Astashyn et al. 2024). Bacterial contigs were then combined into one reference contaminant file. The BBMap v39.08 script *seal.sh* with parameter “ambig=all” and the reference contaminant file was used to decontaminate nanopore reads of bacterial sequences (Bushnell 2014). Hifiasm v0.25.0-r726 with the parameter “--ont" was used to assemble the decontaminated nanopore reads (Cheng et al. 2021). Polypolish v0.6.0 was used to polish the genome assembly (Wick and Holt 2022). To do so, BWA-mem v0.7.19-r1273 was used to map short Illumina reads to the Hifiasm assembly, and the Polypolish command *filter* was used to filter alignments based on insert size (Li and Durbin 2009). The presence of the telomeric repeat CCCTAA was visually examined at the ends of long contigs, and confirmed using the Seqtk v1.4-r122 command *telo* (Li 2025).

### Repetitive DNA analysis

Earl Grey v7.3 with the default parameters was used to identify and classify repetitive DNA regions in the genome assembly (Baril et al. 2024). RepeatModeler v2.0.9 was used for *de novo* repeat family discovery, and RepeatMasker v4.2.4 was used to annotate and mask repetitive sequences throughout the assembly (Flynn et al. 2020). The databases Dfam v4.0 and MycoMobilome v1.1 were used to support the classification of TEs by the Earl Grey pipeline (Storer et al. 2021; Baril and Croll 2026). Earl Grey produced a masked genome and estimated the Kimura 2-parameter divergence of TE copies. The BEDtools v.2.31.1 command *getfasta* was used to extract copies of predicted TEs (Quinlan and Hall 2010). The seqkit v2.13.0 command *translate* was used to translate TE copies in all six frames (Shen et al. 2016). InterProScan v5.69-101.0 was then used to search for conserved domains in TE copies against the databases CDD v3.20 and Pfam v37.0 (Jones et al. 2014). The RIPper webserver was used to predict regions affected by repeat-induced point (RIP) mutations using a sliding window of 1 kb and step size of 500 kb (Van Wyk et al. 2019). Windows with product index value (TpA/ApT)≥1.1, substrate index value (CpA+TpG)/(ApC+GpT)≤0.75, and composite index value (TpA/ApT) – ([CpA+TpG]/[ApC+GpT])≥0 were considered to be affected by RIP. Here, NpN denotes two adjacent nucleotides joined by a phosphodiester linkage (*p*).

### Gene prediction and functional annotation

BRAKER v3.0.8 was used to annotate genes using as evidence RNA-seq data and protein sequences from different fungal species. RNA-seq reads of *A. anomala* isolate OR1 were obtained from the NCBI SRA database (SRR24417797-SRR24417799) (Cohen et al. 2024). Fastp v1.0.1 with the parameter “--length_required 40” and STAR v2.7.11b with parameters “--alignIntronMax 2000 --outSAMstrandField intronMotif” were used to trim and map RNA-seq reads (Dobin et al. 2013; Chen et al. 2018). Protein sequences from *Magnaporthe oryzae* (GCF_000002495.2), *Verticillium dahliae* (GCF_000150675.1), *Neurospora crassa* (GCF_000182925.2), *Pestalotiopsis fici* (GCF_000516985.1), *Cryphonectria parasitica* (GCF_011745365.1), *Metarhizium brunneum* (GCF_013426205.1), *Colletotrichum lupini* (GCF_023278565.1), *Diaporthe amygdali* (GCF_026229845.1), *Sordaria macrospora* (GCF_033870435.1), and *A. anomala* OR1 (GCA_038502455.1) were used in the BRAKER pipeline to assist with gene prediction.

StringTie v2.1.1 was used to assemble transcripts from the mapped RNA-seq reads (Pertea et al. 2015). ORFfinder v0.4.3 with parameters “-s 0 -outfmt 1 -ml 240” was used to predict open reading frames (ORFs) in the assembled transcripts (Wheeler et al. 2007). GMAP v2021-12-17 was used to map ORFs to the genome assembly and generate a GFF file (Wu and Watanabe 2005). The BEDtools v2.30.0 command *intersect* was used to find and remove ORFs overlapping with existing gene models (Quinlan and Hall 2010). SignalP v5 was used to predict ORFs encoding secreted proteins (Armenteros et al. 2019). ORFs encoding secreted proteins were then added as new gene models to the genome annotation. Helixer v0.3.6 using the *Fungi* lineage was also used to predict gene models, which were then merged with the BRAKER gene models (Holst et al. 2026). BUSCO v5.8.3 with parameters “-m prot -l sordariomycetes” was used to estimate gene completeness.

BLASTp searches (e-value <1e-10) of predicted proteins against the UniProt/Swiss-Prot database downloaded on 03 November 2024 were used to obtain functional annotations for each gene (The UniProt Consortium 2019). Only BLAST hits with query and subject coverage of at least 50%, amino acid identity greater than 35% or e-value < 1e-80 were retained. The PANNZER2 webserver with Positive Predictive Value (PPV) >=4 was used to obtain GO terms for the predicted genes (Törönen et al. 2018). BLASTp searches (e-value <1e-10) of predicted proteins against the MEROPS v12.5 and TCDB v20241021 databases were used to identify and classify genes encoding proteases and transporters, respectively (Rawlings et al. 2014; Saier et al. 2021).

The dbCAN v5.2.7 tool was used to predict genes encoding carbohydrate-active enzymes (CAZymes) (Zheng et al. 2023). The output of dbCAN was filtered and only CAZymes with a non-empty recommended result were retained. Signalp5 and Signalp6 were used to predict secreted proteins (Armenteros et al. 2019; Teufel et al. 2022). Proteins identified by either software were considered secreted. DeepTMHMM v1.0 was used to identify transmembrane domains in the mature secreted proteins (Hallgren et al. 2022). NetGPI v1.1 was used to predict membrane-associated secreted proteins (Gíslason et al. 2021). EffectorP3 was used to predicted genes encoding candidate effectors (Sperschneider et al. 2016). Effector proteins predicted by EffectorP3, without transmembrane domains and not predicted to be membrane-bound, were considered as candidate effectors.

### Phylogenetic tree and comparative genomics

Protein sequences from other species of the Gnomoniaceae were downloaded from NCBI GenBank in August 2026, and BUSCO v5.8.3 with parameters “-m prot -l sordariomycetes” was used to identify BUSCO genes. The output of BUSCO was given as input to BUSCO_Phylogenomics v20260630 with parameters “--max_markers 500 --random_seed 9” to obtain a supermatrix amino acid alignment of 500 BUSCO genes (McGowan 2026). IQ-TREE v3.1.3 with parameters “-m MFP -B 1000 --alrt 1000” and the supermatrix alignment was used to infer a partitioned maximum likelihood phylogenetic tree (Nguyen et al. 2015). NUCmer from MUMmer v4.0.0 were used to perform intraspecies synteny analyses between *A. anomala* isolates OR1 and OR-Ennis-d (Marçais et al. 2018). The python package JCVI v1.4.16 was used to perform interspecies synteny analysis between *A. anomala* and *Discula destructiva* (Tang et al. 2024). The BEDtools v2.31.1 commands *complement* and *closest* were used to identify intergenic regions and assign them to adjacent genes (Quinlan and Hall 2010). The R package hexbin v1.28.5 was used to visualize intergenic sizes density for all predicted genes (Carr et al. 2026). The R function *phyper* was used to perform enrichment analyses of different functional gene categories within groups of genes with different intergenic sizes. All R scripts were run within R v4.4.1.

### Mitochondrial genome assembly

Filtlong v0.3.1 with parameter “--keep_percent 10” was used to select nanopore reads containing the best 10% of bases (Wick 2025). Minimap2 v2.30-r1287 was used to map the selected nanopore reads to mtDNA fragments from the genome of isolate OR1 (Li 2018; Cohen et al. 2024). SAMtools v1.24 was used to extract mapped reads (Li et al. 2009). The seqtk v1.4-r122 command *sample* was used to randomly subset 20% of the mapped reads, thus reducing the estimated coverage of the mt genome to 80x (Li 2025). Flye v2.9.4-b1799 with parameters “--nano-hq --genome-size 460k --iterations 3” was used to assemble the reads into a single contig representing the mitochondrial (mt) genome (Kolmogorov et al. 2019). To polish the mt genome, the BBMap v39.33 tool BBDuk with parameter “k=29” was used to retrieve mtDNA-derived Illumina reads (Bushnell 2014). The seqtk command *sample* was used to down-sample reads to 80x coverage. BWA-MEM v0.7.19-r1273 was used to map the mtDNA-derived short reads to the long read mt genome assembly, and Polypolish v0.6.0 was used to polish the mt genome. Circos v0.69-8 was used to generate a circular representation of the mitochondrial genome (Krzywinski et al. 2009).

### Mitochondrial genome annotation and comparative genomic analyses

The MFannot webserver was used to generate an initial gene annotation of the mt genome assembly (Lang et al. 2023). STAR v2.7.11b with parameters “--outSJfilterOverhangMin 12 12 12 12 --alignIntronMin 100 and --alignIntronMax 20000” was used to map the RNA-seq reads from *A. anomala* isolate OR1 (SRR24417797-SRR24417798) to the mt genome assembly (Dobin et al. 2013; Cohen et al. 2024). IGV v2.18.2 was used to visualize the mapped RNA-seq reads, which were used to manually curate the genes (Robinson et al. 2011). tRNAscan-SE v2.0.12 with parameter “--gencode gcode.othmito” was used to predict tRNA genes (Lowe and Eddy 1997). The tRNAscan-SE webserver was used to infer secondary tRNA structures (Lowe and Chan 2016). The RNAweasel webserver was used to classify introns into group I and group II (Lang et al. 2007). ORFfinder v0.4.3 with parameters “-ml 300 -g 4” was used to search for intronic ORFs. InterProScan v5.69-101.0 with e-value < 0.01 was used to search for conserved domains in the predicted ORFs (Jones et al. 2014). NUCmer from the MUMmer package v4.0.0 with parameters “--maxmatch --nosimplify” was used to identify duplicated mtDNA sequences (Marçais et al. 2018). The Entrez Direct v24.0 command *efetch* was used to download fungal mitochondrial genomes from the NCBI Organelle database (Kans 2013 Apr 23). The R package RSCUcaller v1.0 was used to perform a relative synonymous codon usage analysis (Maździarz et al. 2025). BLASTn with e-value < 1e-30 was used to query the mt genome against the nuclear genome to search for nuclear mitochondrial DNA (NUMT) segments. Only alignments longer than 500 bp were considered for downstream analyses. The BEDtools v2.31.1 command *merge* with parameter “-d 10000” was used to merge putative NUMT loci. NUCmer from MUMmer v4.0.0 was used to generate alignments of the NUMT regions and the mt genome.

## Results

### The genome of *A. anomala* is organized into nine chromosomes

The genome of *A. anomala* isolate OR-Ennis-d was sequenced with Oxford Nanopore sequencing, and 6.5 M long reads were generated (49.5 Gbp) with an average length of 7.5 kb, and a read N50 of 18.3 kb. A preliminary genome assembly resulted in 398 Mb organized into 198 contigs. A total of 98 contigs with total size of 55.4 Mb were identified as bacterial contaminants and used as a database to filter out the 10% of total reads identified as contaminants (Supplementary Table 1). After assembling decontaminated reads, any smaller contigs that did not belong to the nuclear chromosomes were filtered out (Supplementary Table 2). The final assembly comprise 342.4 Mb organized into just 9 contigs. The nine assembled contigs all contain canonical telomeric repeats (CCCTAA_n_) at both ends, suggesting they represent complete chromosomes. Thus, the nine contigs were named Chr1 to Chr9, according to their size from largest to smallest (Fig. 1a). The size of the chromosomes varies from 17.4 Mb to 46.6 Mb and are mostly composed of repetitive DNA, as described in a later subsection. At 344 Mb, the *A. anomala* assembly is considerably larger than the genomes of other Gnomoniaceae species included in the comparison, which range from 38.1 to 55.5 Mb (Fig. 1b).

**Fig. 1.**
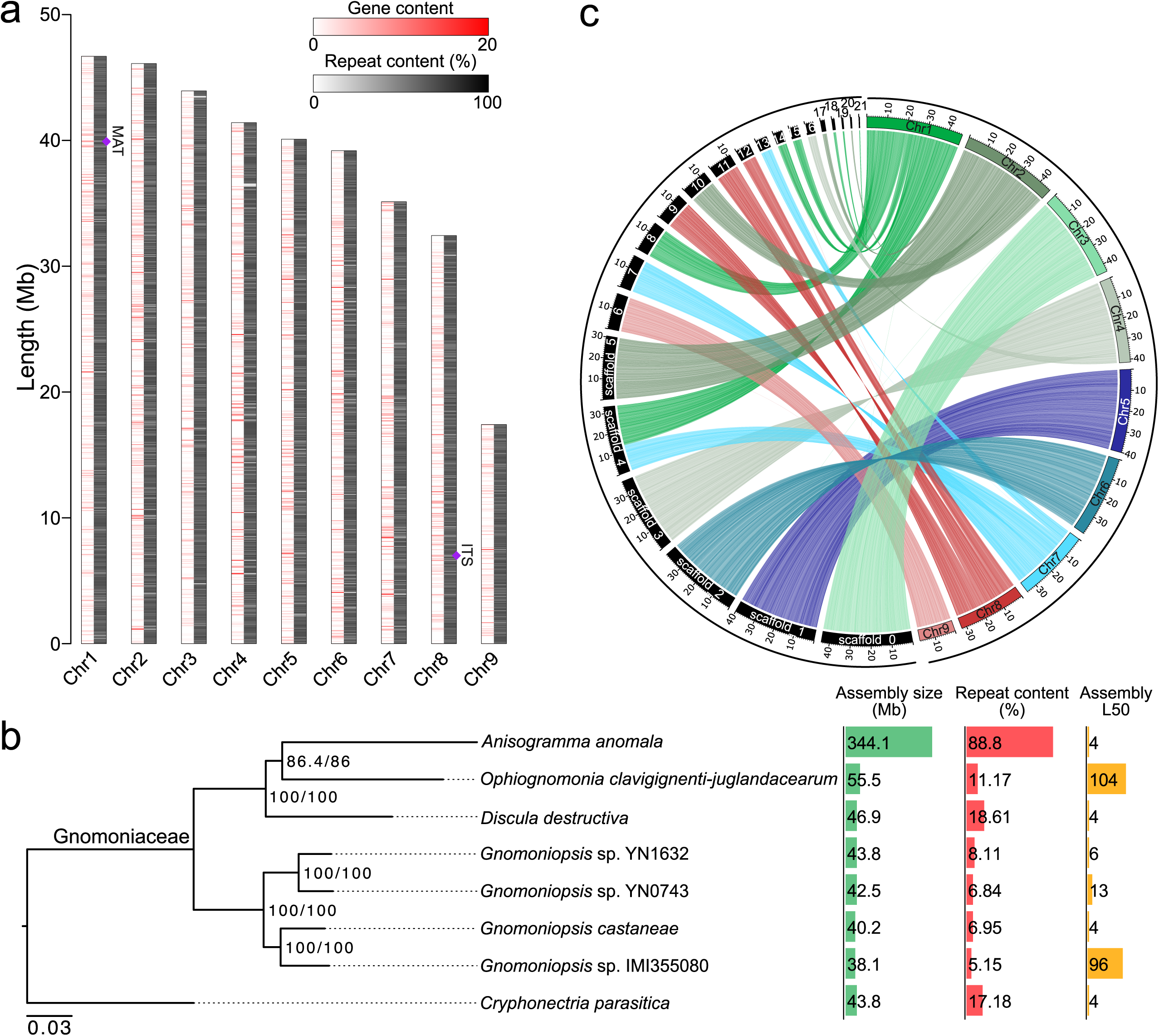
The large genome of *A. anomala* is organized in nine chromosomes. (a) Graphical representation of the predicted chromosomes of *Anisogramma anomala* isolate OR-Ennis-d. Each chromosome is represented by two tracks showing gene content and repetitive DNA content based on a sliding window of 50 kb. Locations of the mating-type locus and ribosomal DNA are indicated. (b) Partitioned maximum-likelihood phylogeny of *A. anomala* and selected Sordariomycetes species. The tree was inferred from a concatenated supermatrix of 500 single-copy BUSCO proteins shared among all analyzed species. Node labels indicate SH-aLRT support/ultrafast bootstrap (UFBoot) support, calculated from 1,000 replicates each. Branch lengths represent the estimated number of amino-acid substitutions per site. The tree was rooted at *C. parasitica*. (c) Whole genome alignment of the predicted chromosomes of *A. anomala* OR-Ennis-d and the assembled scaffolds of *A. anomala* OR1. Ribbons connect homologous regions, colors indicate OR-Ennis-D chromosome. Only scaffolds longer than 100 kb are shown.

A total of 10,676 genes were predicted in the genome of *A. anomala* OR-Ennis-d, which are sparsely distributed throughout the chromosomes at a density of 31 genes per Mb (Fig. 1a). Homology-based functional annotation of the predicted genes indicated the presence of 364 genes encoding carbohydrate-active enzymes (CAZymes), 130 proteases, 1185 transporters, 29 secondary metabolite gene clusters, and 1114 secreted proteins, of which 450 were classified as candidate effectors (Supplementary Tables 3-7). The number of predicted genes in the genome of OR-Ennis-d is considerably higher than the 9,179 reported for isolate OR1 (Cohen et al. 2024) (Table 1). Indeed, there were 1901 genes in OR-Ennis-d that were not predicted in OR1. Among these genes, 45 are BUSCOs and 267 encode secreted proteins. Consistent with this, OR-Ennis-d has a considerably higher BUSCO completeness of 94.3% compared to the 86.6% estimated for OR1. A whole-genome alignment of OR-Ennis-d with the genome of isolate OR1 resulted in 2756 syntenic blocks with an average size of 124.7 kb and 99.95% nucleotide identity, covering 99.5% and 99.6% of the genomes of OR-Ennis-d and OR1, respectively (Fig. 1c). Only one large-scale structural variation was observed between OR-Ennis-d and OR1. Scaffold_4 of the OR1 genome is composed of fragments of Chr1 and Chr7 from OR-Ennis-d (Fig. 1c). However, visual inspection of mapped reads confirmed that this chromosomal rearrangement is likely a misassembly in the genome of OR1 (Supplementary Fig. 1). These results indicate that the genome of isolate OR-Ennis-d is an improvement over OR1, while both isolates have high genome similarity with limited presence-absence variation.

**Table 1.** Genome assembly and gene prediction statistics of *A. anomala* isolates OR-Ennis-d and OR1.

| Stat | OR-Ennis-d | OR1 |
| --- | --- | --- |
| Assembly size (bp) | 342,407,763 | 342,498,352 |
| Contigs | 9 | 2966 |
| Scaffolds | 9 | 112 |
| L50 | 4 | 5 |
| L90 | 8 | 12 |
| N50 (Mb) | 41.4 | 33.2 |
| N90 (Mb) | 32.4 | 0.07 |
| GC% | 34.5 | 34.5 |
| T2T scaffolds | 9 | 2 |
| Predicted genes | 10674 | 9179 |
| Gene completeness |  |  |
| Complete BUSCOs (%) | 94.3 | 86.6 |
| Complete single-copy BUSCOs (%) | 94.1 | 83.6 |
| Complete duplicated BUSCOs (%) | 0.1 | 3 |
| Fragmented BUSCOs (%) | 1 | 5 |
| Missing BUSCOs (%) | 4.8 | 8.4 |

### The genome of *A. anomala* shows no signature of recent TE proliferation

Analysis of repetitive DNA regions indicated that 88.8% of the genome of *A. anomala* OR-Ennis-d is repetitive, consistent with the isolate OR1 for which repeats comprise 89.4% of its genome (Cohen et al. 2024) (Supplementary Table 8). While DNA transposons cover a small percentage (0.14%) of the OR-Ennis-d genome, retrotransposons represent 81.6% of the total genome size, with long terminal repeat (LTR) elements representing 78.5% of the genome. Nearly all LTR elements were classified into two superfamilies, Copia and Gypsy. These two superfamilies represent 59.3% and 19.1% of the OR-Ennis-d genome, respectively, in agreement with the TE content reported for isolate OR1 (Cohen et al. 2024).

To investigate the relative timing of TE accumulation, each annotated TE copy was compared with the consensus sequence of its corresponding TE family. Sequence divergence was estimated using the Kimura 2-parameter (K2P) model, which accounts for differences in the frequencies of transition and transversion substitutions. Because newly inserted TE copies are expected to be highly similar to their family consensus sequence, low K2P divergence values are interpreted as evidence of relatively recent insertion or proliferation. In contrast, higher K2P divergence values indicate that TE copies have accumulated more substitutions since insertion and are therefore relatively older (Oliveira et al. 2025). The repeat landscape of *A. anomala* showed a unimodal and asymmetric distribution of K2P peaking between 2.6 and 2.4 substitutions per 100 sites (Fig. 2a). The distribution extended gradually toward higher K2P values, representing more divergent TE copies, but declined sharply toward lower K2P values. This pattern suggests that most TE accumulation was in the past, with little evidence for recent proliferation. A similar pattern was observed when individual TE subclasses were analyzed separately (Fig. 2b). This pattern contrasts with repeat-rich fungal genomes that contain abundant low-divergence TE copies (Frantzeskakis et al. 2018; Corre et al. 2025).

**Fig. 2.**
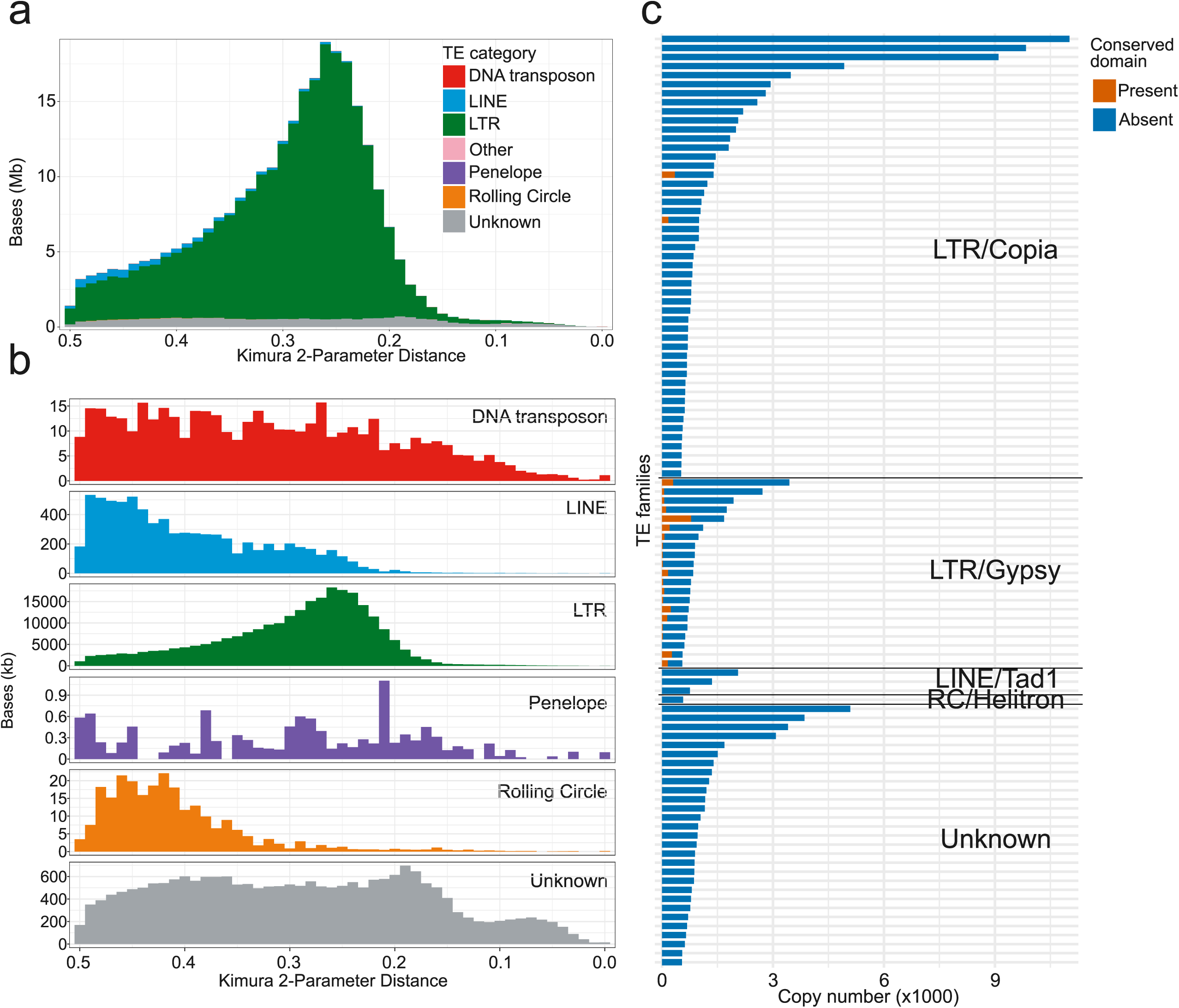
*A. anomala* shows no evidence of recent transposable element (TE) proliferation. (a) Histogram of the pairwise Kimura 2-parameter distance among all TEs in the OR-Ennis-d genome by TE class. Colors indicate TE category. The *x* axis indicates divergence from consensus in Kimura distance, and the *y* axis indicates the accumulated length of TEs with the respective divergence. (b) Histogram of pairwise Kimura 2-parameter distances shown separately for DNA transposons, LINEs, LTR retrotransposons, Penelope-like elements, rolling-circle elements, and unclassified repeats. (c) Copy numbers of the most abundant TE families (>500 copies) in the OR-Ennis-d genome. Horizontal bars represent individual TE families (n = 103 families) and are grouped by TE superfamily or subclass (LTR/Copia, LTR/Gypsy, LINE/Tad1, RC/Helitron, and Unknown). Bar length indicates family copy number. Orange segments denote the number of copies with at least one conserved protein domain (e-value < 0.01) detected in a six-frame translation; blue segments denote copies lacking a detected conserved domain.

An analysis of Repeat-Induced Point (RIP) mutations revealed that 49.7% of the OR-Ennis-d genome is predicted to be affected by RIP. Regions affected by RIP are on average 4.1 kb in length and are distributed throughout all the chromosomes (Supplementary Fig. 2). Consistent with RIP targeting repetitive regions, 51.2% of TE copies overlap with 500-bp windows predicted to be affected by RIP. Given this targeting of TEs by RIP, we next investigated to what extent TE copies in the genome are predicted to be mobilizable or functional. The 103 most abundant TE families with copy number greater than 500, totaling 267 Mb (77%) of the genome, were assessed for conserved TE domains. Surprisingly, only 27 of 103 TE families have at least one copy with a conserved TE domain. Of a total of 154,148 copies of the 103 families, only 3,368 copies (2.2%) have a conserved domain (Fig. 2c). A similar percentage was obtained when considering all 353 TE families identified with at least 10 copies in the genome. Out of the 190,250 TE copies of these 353 families, 4,705 (2.4%) have a conserved domain. This indicates that most TEs in *A. anomala* accumulated mutations that disrupted protein domains required for their autonomous mobility.

### The genome of *A. anomala* is extensively compartmentalized

Despite the fact that 88% of the OR-Ennis-d genome is comprised of TEs, these elements are not evenly distributed throughout the genome. Many intergenic regions are short and absent of TEs, whereas others are long and almost entirely composed of TEs. This organization resembles an extreme form of the architecture of fungal genomes thought to possess a ‘two-speed genome’ model of evolution (Frantzeskakis et al. 2019). A bimodal distribution of GC-content reported for isolate OR1 also suggested that *A. anomala* has a compartmentalized genome (Cohen et al. 2024).

Genes of OR-Ennis-d could be organized into four groups (A, B, C, and D) based on the size of their adjacent intergenic regions (Fig. 3, a and b). Group A, representing genes in gene-dense regions, contains 50.8% of the total genes. These genes are characterized by short, TE-poor flanking intergenic regions, with an average upstream length of 1014 bp and an average downstream length of 922 bp (Fig. 3, a and c). Group B, representing genes with a short intergenic region upstream and a large region downstream, contains 19.9% of the total genes. The TE-poor upstream intergenic regions for group B are an average length of 970 bp, and TE-rich downstream intergenic regions have an average length of 95 kb. Similarly, group C, representing genes with a long upstream and short downstream intergenic region, contains 19.3% of the genes. The TE-rich upstream intergenic regions of group C have an average length of 109 kb, and TE-poor downstream intergenic regions have an average length of 922 bp. The remaining 9.9% of genes belong to group D. Genes in group D are flanked on both ends by long, TE-rich intergenic regions with an average length of 99 kb. As expected, LTR elements were most abundant in TE-rich intergenic regions (Fig. 3c). TEs appeared to be more abundant toward the center of intergenic regions, and less abundant next to the start or end coordinates of the genes. Moreover, no abrupt change in TE composition was observed throughout the length of intergenic regions (Fig. 3c).

**Fig. 3.**
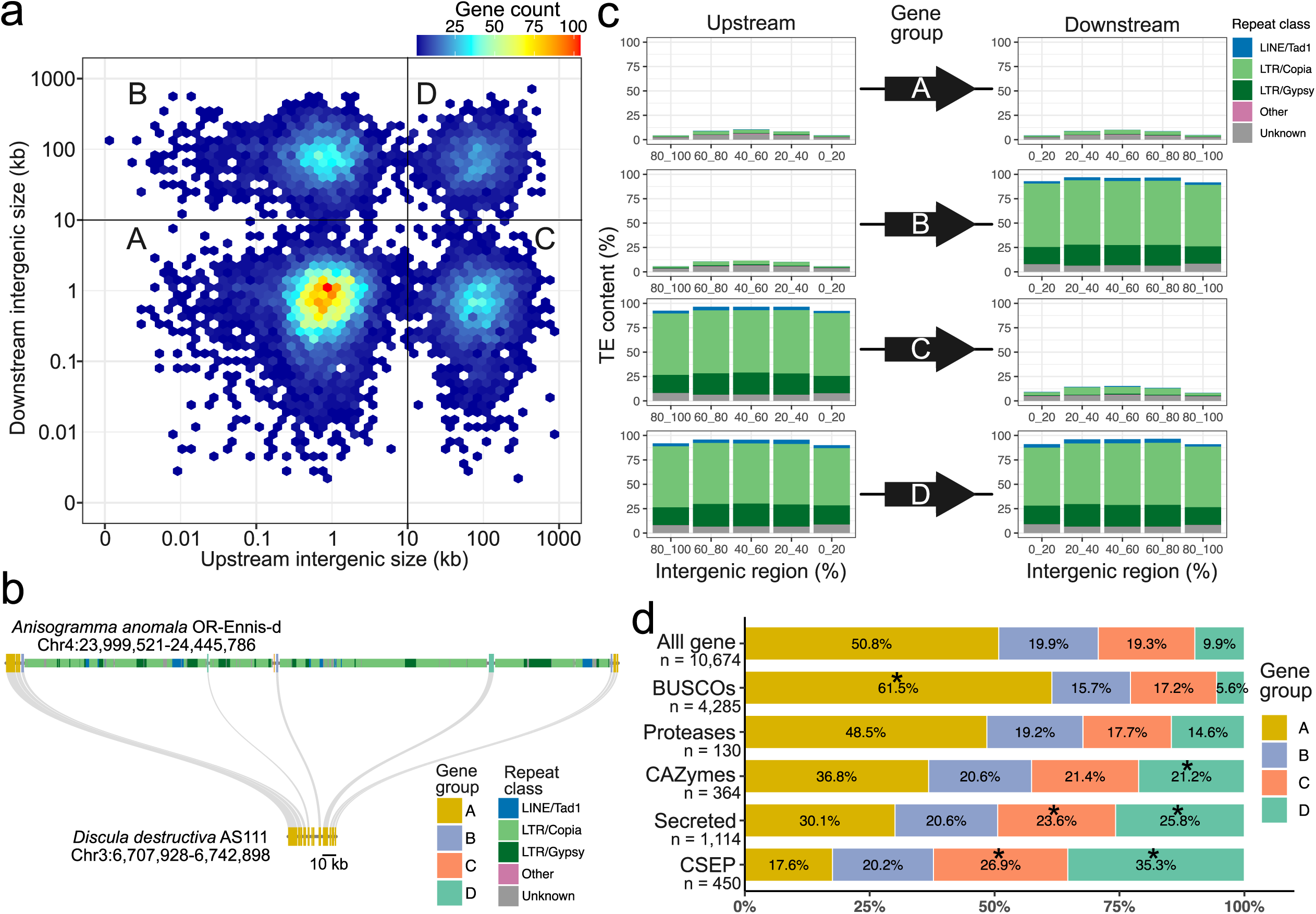
The genome of *A. anomala* is extensively compartmentalized. (A) Heat map showing the distribution of downstream and upstream intergenic sizes for all genes in the genome of isolate OR-Ennis-d. Genes were classified into four groups based on their intergenic size (greater than or less than 10 kb). (b) Local synteny between a 446 kb representative region of chromosome 4 of *A. anomala* and a 35 kb region of chromosome 3 of *Discula destructiva*. Genes are represented as thick rectangles and color-coded based on their intergenic sizes, as shown in (a). Repetitive DNA is also shown as narrow rectangles and color-coded based on their classification. (c) Overall percentage of repeats within intergenic regions of genes from the four main groups. For each gene group A, B, C, and D, the overall transposable element content in up- and downstream intergenic regions is shown as stacked barplots. Intergenic regions were split into five equal parts corresponding to 20% parts, such that 0_20 represents the first 20% of the intergenic region size either upstream or downstream. The stacked barplots represent TE content within each 20% interval as shown on the x-axis. The TE class “Other” represent the sum of the percentages of TE classes that cover less than 0.2% of all the segments of intergenic regions. (d) Percentage of genes from different gene categories that belong to groups A, B, C, and C, as shown in (a). Asterisks indicate significant enrichment of functional gene categories based on hypergeometric tests (p-value < 0.05).

Different functional categories of genes varied significantly in their distribution among the four intergenic-region groups (Fig. 3d, Supplementary Fig. 3). BUSCO genes were enriched in gene-dense group A (P = 4.52e-73), with 61.5% of BUSCO genes assigned to this group, compared with 50.8% of all predicted genes. Conversely, only 5.6% of BUSCO genes occurred in the gene-sparse group D, despite this group containing 9.9% of all genes. In contrast, putative pathogenicity-related genes were preferentially associated with groups C or D. CAZymes were enriched in group D (21.2%; P = 6.26e-11), as were genes encoding secreted proteins (25.8%; P = 4.55e-60) and candidate secreted effector proteins (CSEPs; 35.3%; P = 4.13e-51). Enrichment of genes encoding secreted proteins (23.6%; P = 1.05e-4) and CSEPs (26.9%; P = 4.1e-5) was also observed in group C, although to a lesser extent than group D. No tested gene category was enriched in group B.

### The 462-kb mitochondrial genome of *A. anomala* is one of the largest among fungi

In addition to a greatly expanded nuclear genome, *A. anomala* also has a large mitochondrial (mt) genome. The mt genome of OR-Ennis-d was assembled into a single circular contig of 462 kb with a GC content of 32.9% (Fig. 4a). A total of 167 genes were predicted in the mt genome of *A. anomala*, all having the same transcription orientation. Of these, 14 conserved mitochondrial genes encode proteins involved in the electron transport chain and oxidative phosphorylation, 41 encode mt tRNAs, and two encode mt rRNAs (*rns* and *rnl*). The putative ribosomal protein S3 (*rps3*) gene is located within an intron of *rnl* (Table 2). There are also 109 ORFs within introns that encode proteins similar to homing endonucleases of the LAGLIDADG and GIY-YIG families, or reverse transcriptases (RTs).

**Fig. 4.**
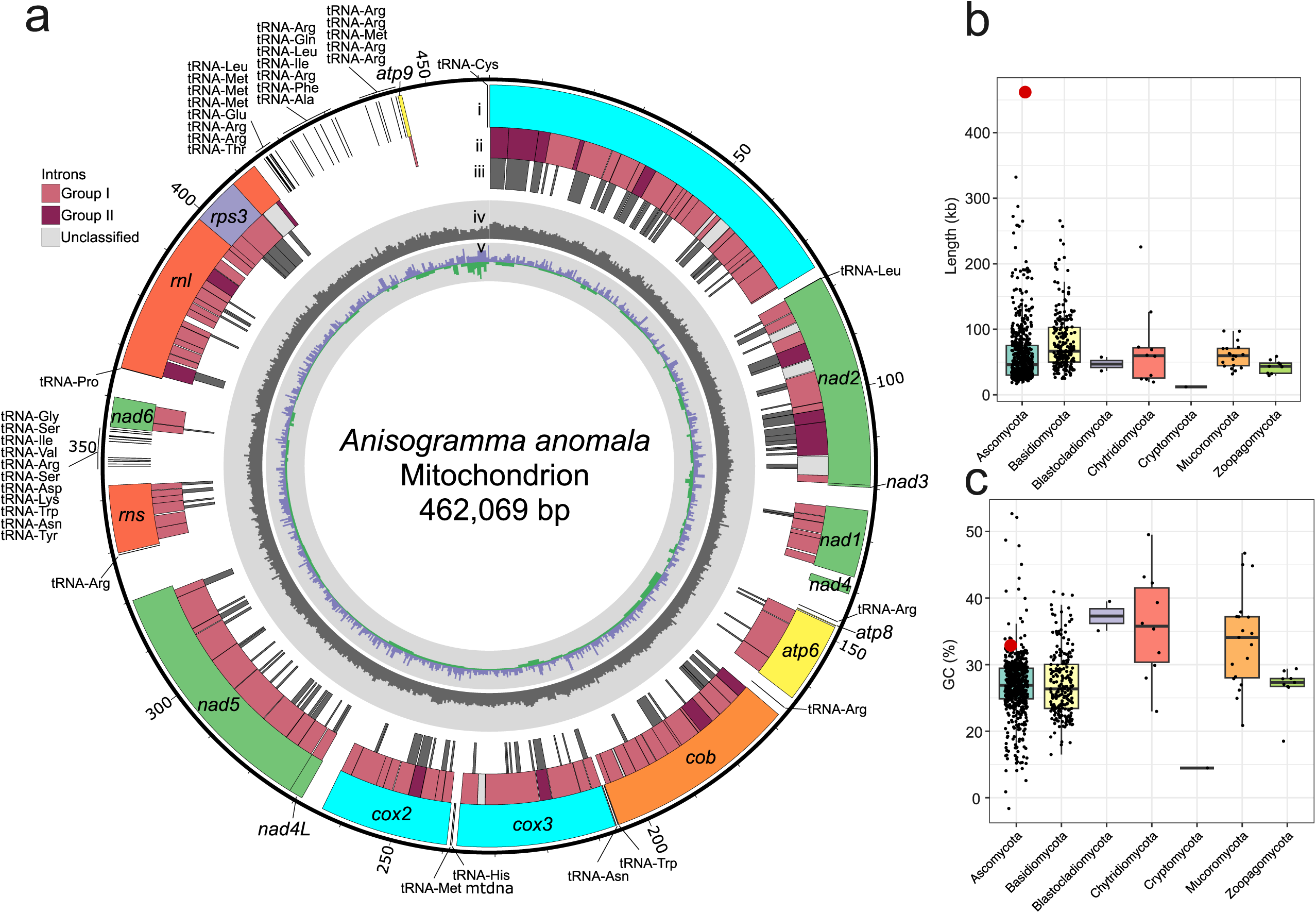
*A. anomala* has one of the largest fungal mitochondrial genomes. (a) Circular representation of the mitochondrial genome of *A. anomala* OR-Ennis-d. Tracks from outside in: (i) Predicted genes commonly found in fungal mt genomes. These genes include genes encoding the subunits of complex I (*nad1*, *nad2*, *nad3*, *nad4*, *nad4L*, *nad5* and *nad6*; in green), complex III (*cob*; in orange), complex IV (*cox1*, *cox2* and *cox3*; in cyan), the ATP-synthase complex (*atp6*, *atp8*and *atp9*; in yellow), the small and large ribosomal subunits (*rns* and *rnl*; in red), the ribosomal protein S3 (*rps3*; in purple), and a set of tRNAs. (ii) Location of mitochondrial introns, classified into Group 1, Group 2, and unclassified. (iii) Predicted open reading frames. (iv) Histogram showing the percent GC content (min = 0%; max = 100%). (v) Histogram showing GC skew as calculated by the formula (G-C)/(G+C). (b) Size distribution and (c) GC content of 841 fungal mitochondrial genomes organized by phylum. Values for *A. anomala* are indicated with larger points colored red.

**Table 2.**
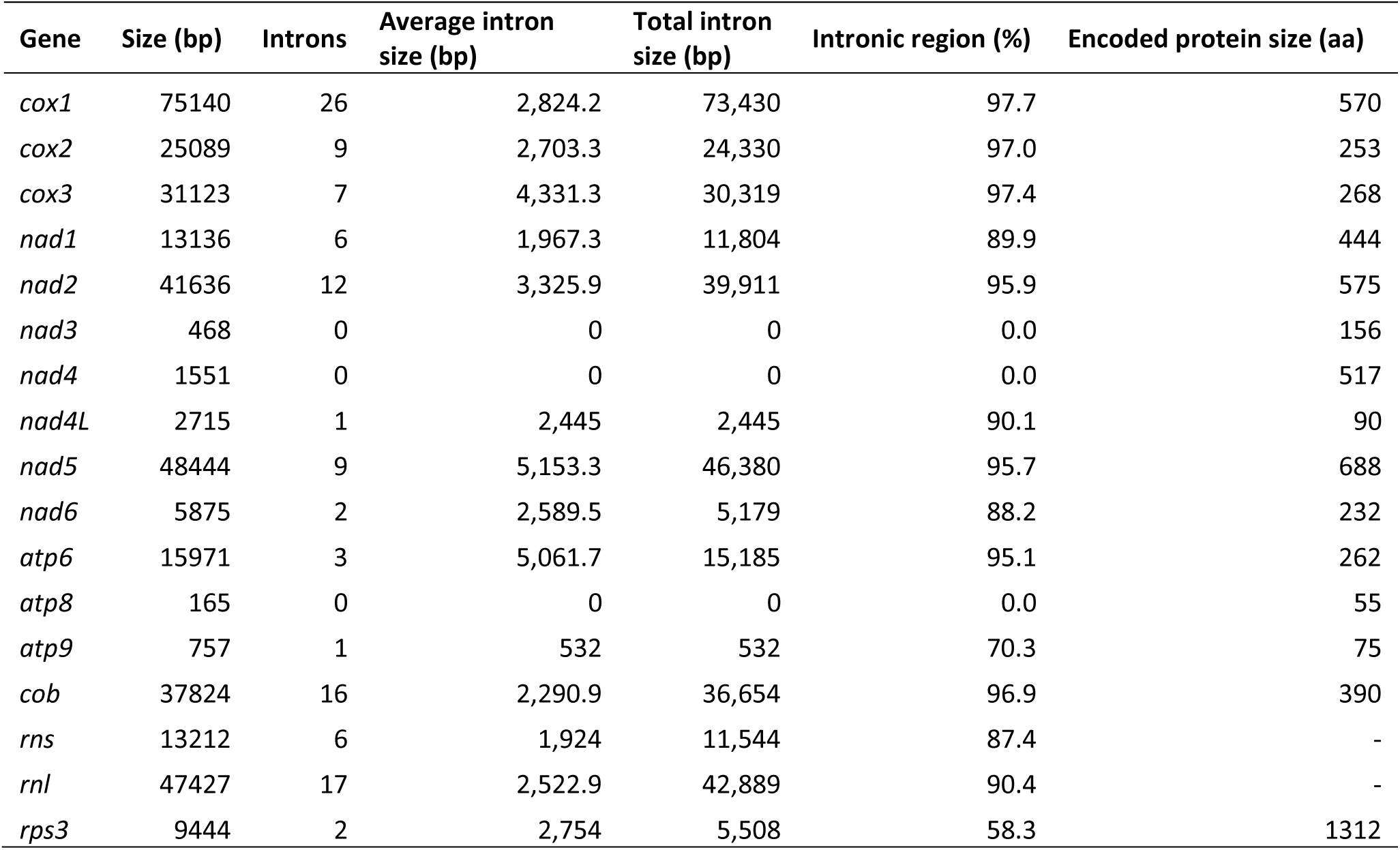
Overall statistics of the core mitochondrial genes of *A. anomala* isolate OR-Ennis-d.

To investigate how the size of the mt genome of *A. anomala* compares to other fungi, we compared the OR-Ennis-d mt genome to 840 representative fungal mt genomes obtained from the NCBI database. Most of these mt genomes were from species of the Ascomycota (n = 584) or Basidiomycota (n = 216) but also included between one and 19 mt genomes from other fungal phyla (Supplementary Table 9). The average size of all fungal mt genomes was 66.5 kb, with a median value of 53.4 kb and standard deviation (SD) of 47.5 kb (Fig. 4b). Ascomycota have overall smaller mt genomes (average = 61.8 kb, median = 45.6 kb, SD = 48.2 kb) compared to Basidiomycota (average = 81.3 kb, median = 66.2 kb, SD = 44.6 kb). Despite that, the three largest mt genomes were from Ascomycota. *A. anomala* has the largest mt genome (461.8 kb), followed by the powdery mildew pathogen *Golovinomyces cichoracearum* (332 kb) and the Ectomycorrhizal fungus *Tuber calosporum* (287 kb) (Li et al. 2020; Zaccaron and Stergiopoulos 2021). The GC content of the mt genome of *A. anomala* was also high relative to other fungi (Fig. 4c). The average GC content of all fungal mt genomes was 27.2%, with Ascomycetes and Basidiomycetes having similar distributions of GC content, with averages of 26.8% and 27.1%, respectively, which are lower than the 32.9% of the mt genome of *A. anomala* (Fig. 4c). These results highlight the variation in size of fungal mt genomes and demonstrate that *A. anomala* has one of the largest mt genomes in the fungal kingdom.

### Intron content explains the large mt genome of *A. anomala*

Unlike the nuclear genome, where TEs and other repetitive DNA explain the expanded genome size, the expansion of the mt genome of *A. anomala* is due to the presence of introns (Supplementary Fig. 4). Introns represent 73.7% of the mt genome. A total of 117 introns were identified on the OR-Ennis-d mt genome, ranging in size from 304 bp to 11.8 kb, and having an average length of 2.9 kb (Supplementary Table 10). Some of these introns are predicted to contain other ORFs. We identified 73 introns (62%) containing ORFs encoding proteins with conserved domains, while 44 introns (38%) are absent of ORFs.

Of the 117 total introns, 110 could be classified as group I or group II introns (Supplementary Table 10). Group I introns are considerably more common than group II introns in the OR-Ennis-d mt genome. Specifically, 93 introns were classified as group I, 16 as group II, and one is a hybrid group I/II intron. This hybrid intron is intron number 15 of the *rnl* gene, and contains the *rps3* gene. The presence of the *rps3* gene within an *rnl* intron is commonly observed in fungi (Korovesi et al. 2018). Interestingly, *rps3* itself contains two introns, a group I (subgroup IC2) intron and a group II intron, both supported by RNA-seq data (Supplementary Fig. 5). Moreover, the predicted *rps3* gene in *A. anomala* is fused with a homing endonuclease gene of the LAGLIDADG family, which has been reported previously in the fungus *Ophiostoma novo-ulmi* (Supplementary Fig. 5) (Sethuraman et al. 2009).

The gene *cox1* is the richest in introns, containing a total of 26 introns. The gene *cob* is also rich in introns, containing a total of 16. Interestingly, intron 9 of *cob* is adjacent to codon 143 (Supplementary Fig. 6). Mutation of this codon is associated with tolerance to quinone outside inhibitor (QOI) fungicides (Fernández-Ortuño et al. 2008). However, the presence of an intron adjacent to this codon is associated with the inability to acquire the mutation p.G143A conferring QOI resistance (Luo et al. 2010).

### *Anisogramma anomala* has an expanded repertoire of mitochondrial tRNA-Arg genes

Examination of the OR-Ennis-d mt genome revealed that *A. anomola* carries an unusually high number of tRNA genes. To investigate how the number of tRNA genes in the mt genome of *A. anomala* compares to other fungi, we compared it to the tRNA gene content of 202 mt genomes from other species of the Sordariomycetes (Supplementary Table 11). Based on the 203 mt genomes analyzed, the average number of tRNA genes was found to be 24.2 with a median of 27 tRNA genes (Fig. 5a). The mt genome of *A. anomala* had the largest repertoire of tRNA genes, with 41, followed by *Nemania diffusa* (n = 36), *Apiospora phragmitis* (n = 34), and *Xylaria hypoxylon* (n = 34). The expanded tRNA gene content in *A. anomala* is mostly due to a large number of tRNA-Arg genes (n = 12), each encoding for one of several anticodons associated with the arginine residue (Fig. 5b, Supplementary Table 12). In comparison, the second largest complement of tRNA-Arg genes was found in the *A. phragmitis* mt genome (n = 5), while the median number of tRNA-Arg genes among all 203 mt genomes was two. The number of tRNA-Met (n = 5), tRNA-Leu (n = 3), and tRNA-Ile (n = 2) genes in the mt genome of *A. anomala* is also high compared to other fungal species (Fig. 5b). A self-alignment of the mt genome of *A. anomala* with itself indicated that many tRNA-Arg and tRNA-Met genes are encoded within short interspersed duplications varying from 67 bp to 595 bp (Supplementary Fig. 4). Thus, small mtDNA duplication events likely contributed to enlarge the arsenal of tRNA genes in *A. anomala*.

**Fig. 5.**
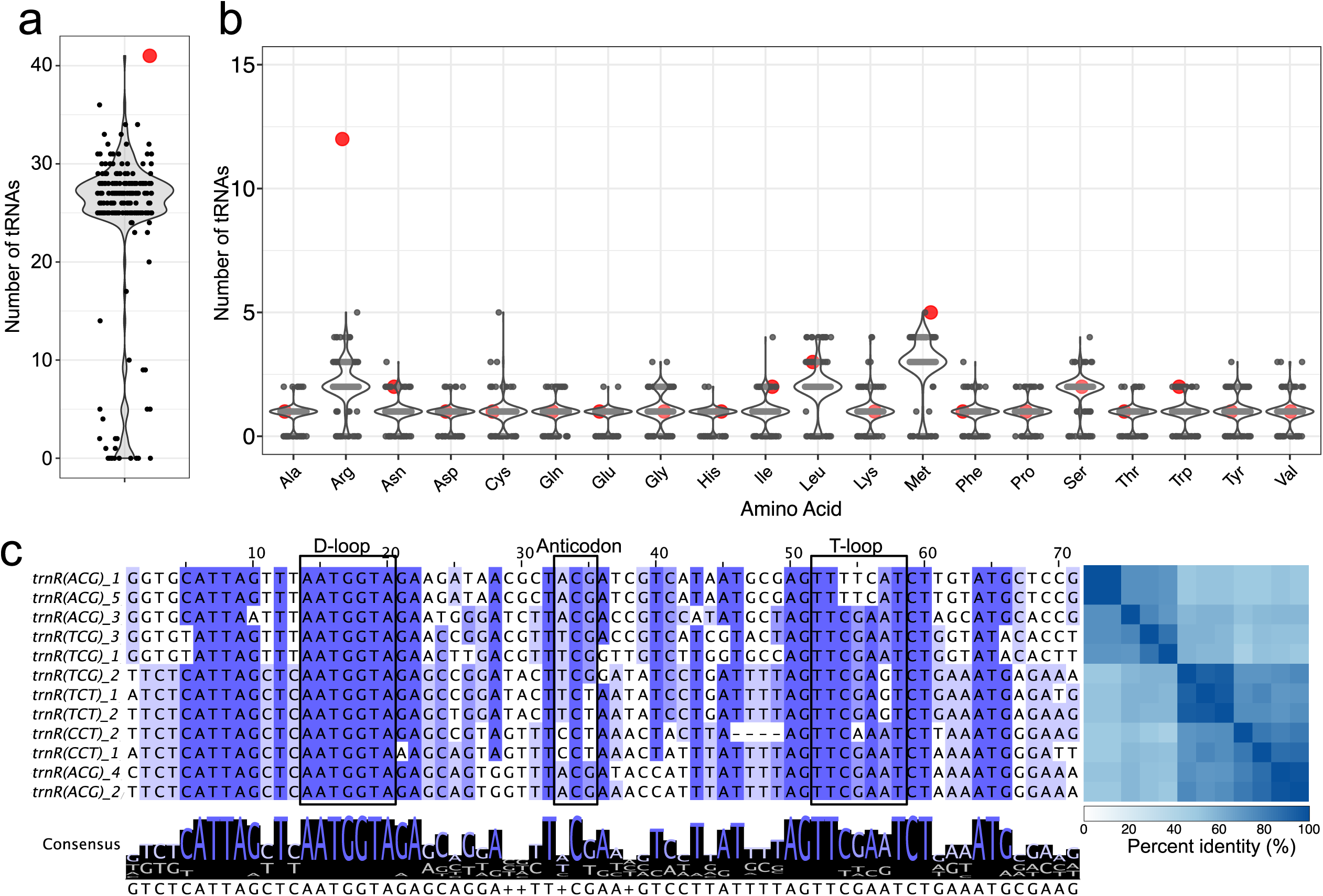
The mitochondrial genome of *A. anomala* is enriched in tRNA-Arg genes. (a) Total number of predicted tRNA genes in 203 mt genomes of species of the Sordariomycetes. In this violin plot, each point represents a mt genome. Value for *A. anomala* is indicated with a larger red point. (b) Number of predicted tRNA genes recognizing the respective amino acid. In these violin plots, each point represents one of the 203 mt genomes analyzed. Values for *A. anomala* are indicated with large red points. (c) Multiple sequence alignment of tRNA-Arg genes in the mitochondrial genome of *Anisogramma anomala*. Conserved sites are indicated with dark blue. The consensus sequence is shown below the alignment. Regions predicted to form the D-loop and the T-loop of the secondary tRNA structures are indicated, as well as the anticodons. A pairwise percent identity matrix is shown on the right.

The predicted secondary structures of all 12 mitochondrial tRNA-Arg genes from *A. anomala* conform with the typical cloverleaf structure of tRNAs, formed by an acceptor stem, the D and T arms, and the anticodon arm (Supplementary Fig. 7). Moreover, the sequences of the 12 tRNA-Arg were conserved, particularly the D-loop sequence that was comprised of the heptanucleotide 5’-AATGGTA-3’ (Fig. 5c). This suggests that all 12 tRNA-Arg genes in *A. anomala* are likely to be functional.

Despite the high abundance of tRNA-Arg genes, no abnormal usage of arginine-coding codons was observed in the mt genome of *A. anomala*. Specifically, the percentage of arginine residues observed across all 14 core protein coding genes was 2.3%, which is comparable to other fungi that vary from 1.9% to 3.0% (Supplementary Fig. 8a). Similar patterns were observed when analyzing each gene individually (Supplementary Fig. 8b). Bias toward the arginine AGA codon (relative synonymous codon usage [RSCU] = 3.9) was observed, whereas the other arginine codons were underused (RSCU ≤ 0.83) (Supplementary Fig. 8c). However, only two tRNA genes were predicted with anticodons matching to AGA, whereas five tRNAs matched to codon CGT (RSCU = 0.83), and three tRNAs matched to codon CGA (RSCU = 0.44), which are underused codons (Supplementary Fig. 8d). Thus, the large arsenal of tRNA-Arg in the mt genome of *A. anomala* is unlikely due to high translation demand of arginine codons.

### The *A. anomala* nuclear genome contains large mitochondrial DNA insertions

Inspection of the *A. anomala* nuclear genome revealed several regions characterized by low TE density and an almost complete absence of predicted genes. These regions showed substantial similarity to the mt genome, resembling nuclear mitochondrial DNA segments (NUMTs) (Lopez et al. 1994). To identify NUMTs, the *A. anomala* mt genome was queried against the nuclear genome assembly using BLASTn. A total of 131 alignments with an e-value < 1e-30 and an alignment length greater than 500 bp were retained. These alignments had a mean nucleotide identity to the mt genome of 89.8% and collectively covered 616.9 kb of nuclear sequence. BLASTn hits occurring within 10 kb in the nuclear genome were merged, resulting in 20 putative NUMT loci. Individual NUMT loci range in size from 522 bp (NUMT10) to 220.1 kb (NUMT13).

The NUMTs are distributed across eight of the nine nuclear chromosomes and show no clear enrichment in particular chromosomal regions (Fig. 6a). Mapped long nanopore reads span the junctions between NUMTs and their flanking nuclear sequence, further supporting their placement in the nuclear assembly instead of as assembly errors (Supplementary Fig. 9). To assess conservation of NUMTs in other *A. anomala* isolates, 1-kb sequences spanning each NUMT junction were extracted from the OR-Ennis-d assembly and aligned to the OR1 reference genome. All 40 boundary-spanning queries aligned to the OR1 genome assembly with >97% query coverage and >99% nucleotide identity. This result supports conservation of the NUMT insertion junctions in the genome of isolate OR1. Collectively, the NUMTs contain sequences homologous to 74.7% of the *A. anomala* mt genome, including fragments of 11 core mt protein-coding genes (*atp6*, *atp8*, *cob*, *cox1*–*cox3*, and *nad1*–*nad5*), both mt ribosomal RNA genes (*rnl* and *rns*), and 28 mt tRNA genes (Fig. 6b). The mt genomic region containing *nad4* is represented repeatedly among the nuclear insertions. Five large *nad4*-derived fragments occur across NUMT5, NUMT11, NUMT13, and NUMT16. However, only the *nad4* copy in NUMT11 retains a near-intact open reading frame, encoding a predicted protein with 96.5% amino-acid identity to the mitochondrial nad4. The remaining four *nad4* copies contain in-frame stop codons.

**Fig. 6.**
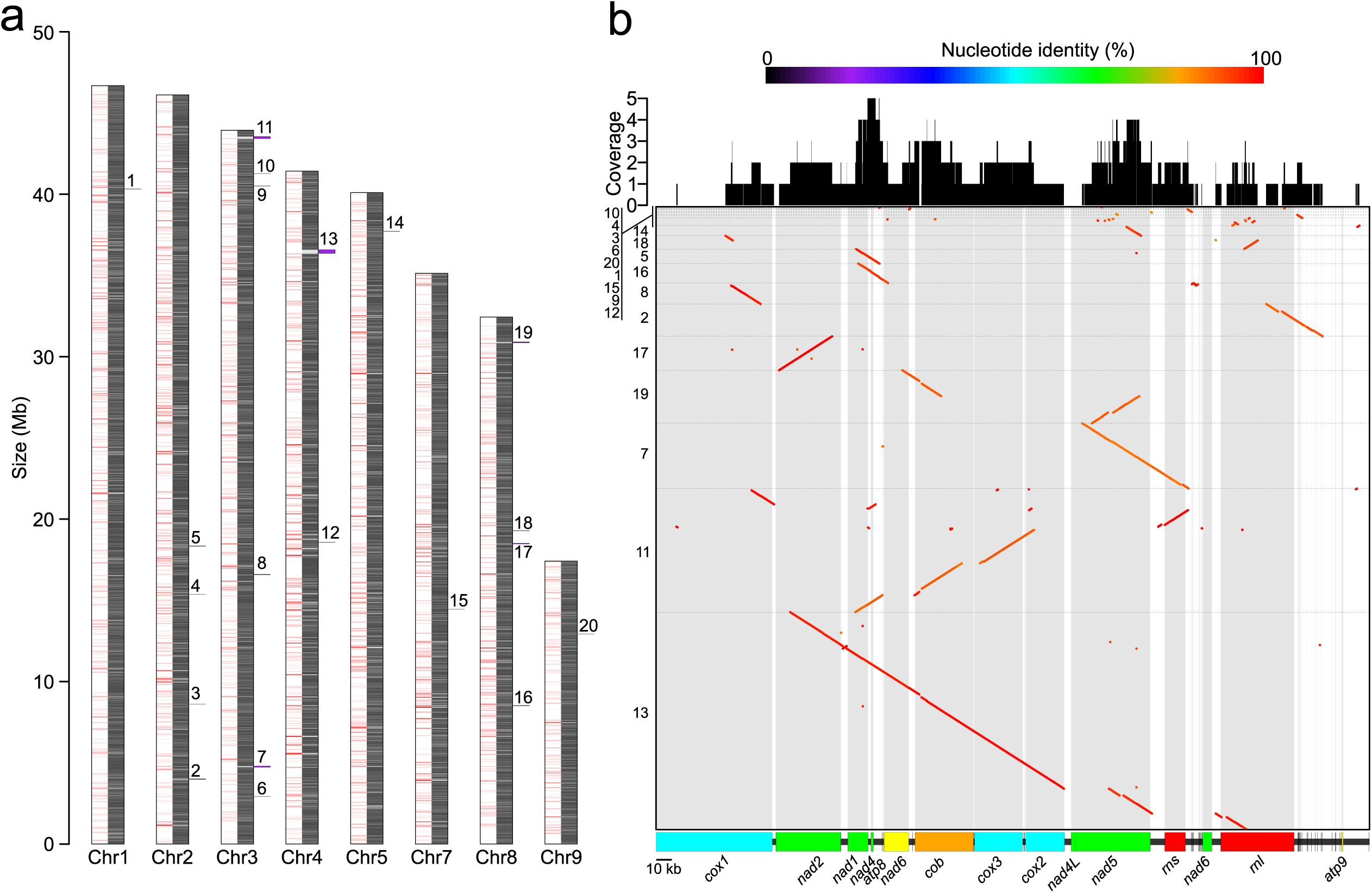
The genome of *A. anomala* contains long nuclear-mitochondrial segments (NUMTs). (a) Location and size of 20 NUMTs throughout the predicted chromosomes of OR-Ennis-d. Each chromosome is represented by two tracks showing gene content (left) and repetitive DNA content (right) based on a sliding window of 50 kb. The positions of NUMTs are labeled 1 to 20 based on their occurrence. (b) Dot plot alignment of the mitochondrial (mt) genome of the OR-Ennis-d mt genome (x axis) with the predicted NUMTs (y-axis). NUMTs are numbered as in panel (a). The mt genome is shown on the x-axis, where rectangles represent core mt genes. Shaded areas highlight the gene space.

Most NUMTs occur within TE-rich genomic regions and are not predicted to disrupt annotated genes (Supplementary Fig. 10). Only four NUMTs have predicted genes within 15 kb of either boundary, and only NUMT3 and NUMT6 occur within 2 kb of an annotated gene. NUMT3 is located 865 bp upstream of gene AAOREND_01475, which encodes a putative magnesium-dependent phosphatase MDP1, whereas NUMT6 is located 1.8 kb upstream of gene AAOREND_02824, which encodes a hypothetical protein. Three gene models were predicted within NUMT loci (AAOREND_02866, AAOREND_05272, and AAOREND_10251). However, these encode hypothetical proteins or proteins with similarity to homing endonucleases.

## Discussion

We generated a chromosome-scale reference genome for *A. anomala*, the causal agent of Eastern filbert blight, the most economically important disease of hazelnut in the United States. One of the most prominent features of the *A. anomala* genome is its extreme compartmentalization, in which approximately half of predicted genes are located in TE-rich, gene-sparse regions enriched in putative pathogenicity-related genes, whereas the remainder are located in TE-poor, gene-rich regions. This architecture is consistent with the genome compartmentalization described in diverse fungal and oomycete pathogens (Dong et al. 2015; Fokkens et al. 2018; Torres et al. 2020; Zaccaron et al. 2022; Wacker et al. 2023). In such genomes, TE-rich regions can promote local genomic plasticity through TE-associated structural variation, including transposition, repeat-mediated recombination, gene duplication or deletion, and genome rearrangement (Frantzeskakis et al. 2019; Torres et al. 2020; Zaccaron and Stergiopoulos 2024). TEs can also affect the regulation of nearby genes, while repeat-induced point mutation (RIP) can increase sequence divergence within duplicated or repeat-associated DNA (Rouxel et al. 2011; Dong et al. 2015). These processes provide the conceptual basis for the ‘two-speed’ genome model, in which genes located in repeat-rich regions may undergo more rapid sequence evolution, gene-content turnover, or regulatory variation than genes in repeat-poor regions (Dong et al. 2015; Torres et al. 2020).

The enrichment of putative pathogenicity-related genes in TE-rich regions raises the possibility that genome compartmentalization contributes to adaptation in the *A. anomala*–hazelnut pathosystem. This possibility is particularly relevant because the extensive deployment of hazelnut cultivars carrying the ‘Gasaway’ resistance locus in the Pacific Northwest likely imposed strong selection on pathogen populations. Isolates capable of causing disease on cultivars carrying ‘Gasaway’-derived resistance have recently been detected, demonstrating that *A. anomala* can overcome this source of resistance (Wiman 2023). TE-rich regions could provide a plastic genomic environment in which pathogenicity-associated genes acquire adaptive mutations. Thus, the compartmentalized architecture of *A. anomala* genome may have facilitated the emergence of resistance-breaking isolates. However, the presence of TE-rich, gene-sparse regions alone does not establish that *A. anomala* exhibits differential evolutionary rates across its genome, nor does it demonstrate a causal role in resistance breakdown. Testing this hypothesis will require comparisons of sequence polymorphism, structural variation, and gene presence-absence variation between the two compartments, particularly among isolates that differ in virulence on ‘Gasaway’-derived cultivars.

The enrichment of genes encoding secreted proteins and candidate secreted effector proteins (CSEPs) in groups C and D, but not in group B, indicates a disproportionate association of these gene categories with long, TE-rich 5′ flanking regions. This asymmetric distribution is unlikely to result from a genome-wide bias toward TE-rich upstream regions, because groups B and C contained similar proportions of all predicted genes. Instead, the pattern supports a category-specific association between CSEP genes and TE-rich upstream sequence. One possibility is that the presence of upstream TEs affects gene regulation by modifying promoter sequences or local chromatin states, thus potentially contributing to expression plasticity of host-interaction genes. Comparative analyses of TE insertion polymorphism, local chromatin features, and infection-associated gene expression will be needed to evaluate this hypothesis.

The extensive compartmentalization of the *A. anomala* genome appears to be a consequence of TE accumulation. Repetitive sequences represent 88.8% of the OR-Ennis-d genome, consistent with the 89.4% repeat content reported for isolate OR1 (Cohen et al. 2024). TE proliferation is a major driver of genome expansion in fungi. Rust fungi (Pucciniales) and entomophthoralean fungi (Entomophthorales) include some of the largest fungal genomes, often approaching or exceeding 1 Gb, and their genome expansions have been attributed largely to TE accumulation (Tobias et al. 2021; Gupta et al. 2023; Stajich et al. 2024). Within the Ascomycota, the obligate powdery mildew fungi likewise typically have unusually large, repeat-rich genomes (Frantzeskakis et al. 2018; Kusch et al. 2022; Roy et al. 2026). These examples illustrate that genome size can increase substantially when TE proliferation exceeds sequence loss. The *A. anomala* genome conforms to this pattern, but its repeat expansion was highly uneven. Instead of uniformly large intergenic distances across the whole genome, transposons accumulated within a subset of intergenic regions.

Despite their abundance, the TE landscape of *A. anomala* OR-Ennis-d shows no evidence of substantial recent proliferation. The aggregate K2P divergence profile is unimodal and sharply depleted at low divergence, instead of showing the low-divergence peak expected after a major recent expansion (Betancourt et al. 2024). In TE landscapes, low K2P divergence from a family consensus sequence generally indicates recently inserted copies, whereas more divergent copies are interpreted as relatively older because they have accumulated mutations after insertion. Thus, the TE landscape of *A. anomala* suggests that nearly all TE accumulation was ancient, followed by inactivation and loss. There is little evidence for recent or active TE accumulation. Moreover, only 2.2% of TE copies from the 103 most abundant TE families retain recognizable conserved protein domains. These findings are consistent with the earlier OR1 genome analysis, in which no intact autonomous LTR retrotransposons were identified (Cohen et al. 2024). Together, these results suggest that the *A. anomala* genome is dominated by degenerated remnants of historical TE amplification events, instead of by a highly active mobilome.

A history of TE amplification and decay could also explain the large percentage of TE copies without clear dinucleotide bias resulting from RIP mutations. As reported herein, only 51.2% of TE copies overlapped regions predicted to be affected by RIP. This indicates that nearly half of the TE copies in the genome of *A. anomala* show no signature of RIP. Although genes implicated in RIP are conserved in the genome of *A. anomala*, RIP could be inefficient or could have affected sequences unevenly across the genome (Cohen et al. 2024). However, another possibility is that TE copies that were previously affected by RIP may have subsequently accumulated additional mutations, deletions, or fragmentation that obscured diagnostic dinucleotide biases. The mixture of RIP-affected and apparently non-RIP-affected TE copies therefore likely reflects a complex history of TE invasion, genome defense, and long-term sequence decay. The homothallic reproductive biology of *A. anomala* may also influence TE persistence by altering effective outcrossing, recombination, and the efficacy of purifying selection (Whittle et al. 2011; Cohen et al. 2024).

In addition to its unusual nuclear genome architecture, *A. anomala* has a large and intron-rich mitochondrial genome. Mitochondrial introns are common in fungi and are major contributors to variation in fungal mitochondrial genome size (Sandor et al. 2018). The 462-kb *A. anomala* mitochondrial genome, which contains 117 introns, is among the largest reported for Ascomycota. One potentially important feature is an intron adjacent to codon 143 of the *cob* gene. The p.G143A substitution is a common mechanism of resistance to quinone outside inhibitor (QoI) fungicides in fungal pathogens (Fernández-Ortuño et al. 2008). However, an intron at this position can prevent the emergence of this substitution because nucleotide changes that generate p.G143A may disrupt intron splicing and yield a nonfunctional cytochrome b transcript or protein. This has been proposed for the leotiomycetes *Botrytis cinerea*, *Monilinia fructicola,* and *Golovinomyces cichoracearum* (Banno et al. 2009; Luo et al. 2010; Zaccaron and Stergiopoulos 2021). The presence of an intron adjacent to codon 143 in *A. anomala* may therefore limit the acquisition of QoI resistance through p.G143A. This possibility is particularly relevant because QoI fungicides are commonly effective for management of Eastern filbert blight (Pscheidt et al. 2017). It will be important to characterize whether all or most *A. anomala* strains contain this intron to inform on management practices.

The unusually large mitochondrial tRNA repertoire of *A. anomala* is most likely the result of local mtDNA duplication. Variation in tRNA copy number can be shaped by codon usage and translational selection (Guerreiro et al. 2026). However, the mt tRNA-Arg expansion in *A. anomala* showed no correspondence with increased arginine codon usage compared to other fungal species. Selection for increased translational demand is therefore unlikely to explain the high copy number of mt tRNA-Arg genes. Nevertheless, all mt tRNA-Arg genes retained canonical predicted cloverleaf structures and conserved sequence features, suggesting that they may remain functional. Whether these duplicated mt tRNA-Arg genes are transcribed, aminoacylated, or contribute to translation of mitochondrial genes remains to be determined.

Another unusual feature of the *A. anomala* genome is the presence of large nuclear mitochondrial DNA segments (NUMTs), which are fragments of mitochondrial DNA transferred to and retained in the nuclear genome (Lopez et al. 1994). NUMTs have been documented across diverse eukaryotes, including fungi such as *Neurospora*, *Trichoderma*, and *Penicillium* species (Hazkani-Covo et al. 2010; Zhang et al. 2017; Li Wan-Chen et al. 2021; Zhang et al. 2021; Requena et al. 2025). Although NUMT insertions that disrupt protein-coding genes have been associated with inherited disease and tumorigenesis in humans (Hazkani-Covo et al. 2010; Wei et al. 2022), their functional and evolutionary consequences in fungi remain poorly understood. In *A. anomala*, most identified NUMTs are in TE-rich regions, and none are predicted to disrupt a protein-coding gene. A similar association between NUMTs and repeat-rich regions has been reported in mammalian genomes (Tsuji et al., 2012). This distribution is consistent with purifying selection acting against deleterious NUMT insertions that disrupt genes, while insertions in TE-rich regions are more likely to persist because they impose smaller fitness costs. The extensive TE-rich compartment of the *A. anomala* genome may therefore provide permissive genomic environments for the long-term retention of NUMTs.

The 20 NUMT loci identified in *A. anomala* collectively spanned at least 616.9 kb of nuclear sequence and contained homologous fragments representing 74.7% of the mitochondrial genome. This amount is substantially larger than the NUMT content reported in an early survey of 20 fungal genomes, in which total NUMT content ranged from 78 bp in *Candida albicans* to 77.1 kb in *Phaeosphaeria nodorum* (Hazkani-Covo et al. 2010). However, this comparison should be interpreted cautiously because NUMT identification is sensitive to mitochondrial and nuclear assembly quality and sequence-similarity thresholds. Nonetheless, the results presented indicate that *A. anomala* harbors an unusually extensive repertoire of NUMTs among currently surveyed fungi. Further comparative analyses will be important to determine whether this feature is exceptional within the Gnomoniaceae or characteristic of exceptionally large, repeat-rich fungal genomes.

In summary, the chromosome-scale *A. anomala* genome reveals an exceptionally repeat-rich and compartmentalized nuclear genome, an expanded intron-rich mitochondrial genome, and extensive mitochondrial-to-nuclear DNA transfer. Historical LTR retrotransposon accumulation appears to have generated long TE-rich intergenic regions that partition conserved core genes from genomic compartments enriched in CAZymes, secreted proteins, and candidate effectors. Although most TE copies appear degenerated and no major recent proliferation was detected, their long-term retention has profoundly shaped genome architecture. Finally, these genomic resources provide a foundation for population genomic surveillance of emerging resistance-breaking isolates, identification of candidate pathogenicity and avirulence genes, and investigation of the evolutionary processes that shape the *A. anomala*–hazelnut interaction.

## Data availability

Genomic data was deposited at NCBI under BioProject PRJNA1507296. The sequenced nanopore and Illumina reads for *A. anomala* OR-Ennis-d were deposited in the NCBI Sequence Read Archive under accessions SRR40127654 and SRR40127653, respectively. The nuclear and mitochondrial genome assemblies of *A. anomala* OR-Ennis-d were deposited in GenBank under the accessions JCCFBL010000000 and PZ891560, respectively.

## Acknowledgments

We thank Dr. Alanna Cohen for providing genomic resources for *A. anomala*. We thank the Center for Quantitative Life Sciences at Oregon State University for proving computational support. We also thank the Oregon Hazelnut Commission for funding and supporting this research. Perplexity AI was used to edit manuscript text for grammar and style.

